# MicroAge Mission: Experimental Design, Hardware Development and Operational Considerations for a Bespoke Culture System to Support Tissue-Engineered Skeletal Muscle Constructs

**DOI:** 10.1101/2025.06.17.660117

**Authors:** Samantha W. Jones, Shahjahan Shigdar, Benjamin Tollitt, Adam Janvier, Fiona Mutter, James R. Henstock, Jessica Ohana, David A. Turner, Christopher McArdle, Gianluca Neri, William Blackler, Georgi Olentsenko, Kai. F. Hoettges, Anne McArdle, Malcolm J. Jackson

## Abstract

Microgravity provides a unique model for understanding accelerated skeletal muscle loss, and potentially a model of muscle ageing, offering insights into the molecular mechanisms underlying reductions in muscle mass and function. During spaceflight, astronauts experience pronounced skeletal muscle atrophy. These effects appear similar to age-related muscle decline on Earth but on a significantly shorter timescale. Despite the incorporation of daily aerobic and resistance exercise on the International Space Station (ISS), countermeasures remain suboptimal, reflecting analogous challenges in exercise efficacy observed in ageing populations.

The MicroAge Mission aimed to exploit microgravity conditions aboard the ISS to determine whether the molecular mechanisms underpinning reduced adaptive responses to contractile activity during ageing are analogous to those induced by spaceflight. The mission also explored proof-of-concept genetic interventions, including overexpression of Heat Shock Protein 10 (HSP10), a mitochondrial chaperone, to mitigate muscle atrophy and functional loss.

To conduct these investigations, a tissue-engineering approach was employed to fabricate human skeletal muscle constructs, which were secured to custom-designed 3D-printed scaffolds. The scaffolds featured integrated microfluidic channels designed to interface with the fluid handling system within the flight hardware. The hardware, developed by Kayser Space Ltd, was specifically designed to interface with the European Space Agency’s (ESA) Kubik incubator located within the Columbus module of the ISS.

This research addresses critical methodological constraints in low Earth orbit (LEO) experimentation, providing a detailed account of pre-flight protocol development, muscle construct biofabrication techniques, and operational considerations. The findings establish a translational framework for future investigations into musculoskeletal degeneration, with implications for therapeutic strategies targeting both terrestrial ageing and astronaut musculoskeletal health.

## 1 Introduction

Microgravity potentially offers a powerful model for understanding accelerated skeletal muscle ageing, providing insights into the molecular mechanisms underlying muscular decline ^1^. During spaceflight, astronauts experience rapid reductions in skeletal muscle mass, with virtually all muscle groups showing a 3-10 % decrease in volume after just 17 days in microgravity ^2–4^. Prolonged exposure (≥ 6 months) exacerbates these effects, leading to significant physiological changes that appear to mirror age-related muscle decline on Earth ^4^. Similar to ageing, microgravity affects specific muscle groups disproportionately, though all fibre types are ultimately compromised ^3,5^. Despite the implementation of daily (∼2 hours/day) aerobic and resistance exercise on the International Space Station (ISS), the preventative effects remain incomplete, echoing the challenges of exercise efficacy in ageing populations ^6,7^.

On Earth, demographic shifts driven by declining fertility and mortality rates have resulted in a growing number of older adults (aged ≥ 65) with relatively poor health and diminished quality of life ^8^. Age-related loss of skeletal muscle mass and function, a major contributor to physical frailty, significantly increases the risk of falls and hospitalisation. By the age of 70, muscle cross-sectional area can decrease by ∼25-30 %, with muscle strength diminishing by ∼30-40 %, accompanied by muscle atrophy and weakened fibres ^9–11^.

In healthy adults, contraction-mediated biochemical responses to exercise drive anabolic changes in skeletal muscle. These adaptive changes are initiated by an acute increase in reactive oxygen species (ROS), which act as redox-signalling molecules under normal physiological conditions ^12–14^. However, ageing is associated with attenuated ROS-mediated adaptive responses, reducing exercise efficacy. This decline is thought to stem from chronically elevated mitochondrial H_2_O_2_ production ^9,15,16^.

The MicroAge Mission aims to leverage microgravity conditions on board the ISS, to investigate whether the molecular mechanisms driving diminished adaptive responses to contractile activity in ageing are the same as those observed during spaceflight. This research holds promise for advancing our understanding of skeletal muscle decline and developing effective interventions for both ageing populations and astronauts ^1^.

Additionally, the mission sought to investigate proof-of-concept genetic intervention strategies for maintaining muscle function, specifically through the overexpression of Heat Shock Protein 10 (HSP10), a mitochondrial chaperone known to offer some protection against loss of maximal tetanic force and loss of muscle cross-sectional area observed in old mice ^17^. The study also evaluated the impact of simulated Earth gravity (1g) using centrifugation, alongside a comprehensive post-flight ground reference experiment (GRE) to generate appropriate ground-based control data.

To carry out these studies, a tissue-engineering strategy was adopted to create human skeletal muscle constructs anchored to custom-designed 3D-printed scaffolds. These scaffolds incorporated microfluidic channels that integrated with a specialised fluid handling system within the flight hardware. The hardware units, developed by Kayser Space Ltd, were engineered to be compatible with the European Space Agency’s (ESA) Kubik incubator, located in the Columbus module of the ISS.

Conducting experiments in low Earth orbit (LEO) presents significant methodological and logistical challenges. While the ISS boasts a rich legacy in cell-based life science research, developed over more than 24 years of continuous operation, considerable variability still exists in the methodologies employed, resource availability, and the standardisation of equipment used in these studies ^18^. To help address these challenges, this paper outlines the development of the MicroAge flight hardware, the pre-flight experimental design, muscle construct biofabrication techniques, and operational considerations for conducting biological research in LEO. A key objective is to share lessons learned, offering a practical framework that other researchers can adapt and build upon for future missions, ultimately contributing to the creation of a more standardised workflow for experiments of this kind.

## 2 Materials, Methods and Hardware Descriptions

### Materials

Human immortalised myoblasts (Lot AB1167: *fascia lata* biopsy, 20 years, male), generated using the MyoLine Platform, were obtained via a Materials Transfer Agreement (MTA) from the Institute of Myology (Paris, France) ^19,20^. Skeletal muscle cell growth medium kits (phenol red-free) were obtained from PromoCell (Dorset, UK).

High glucose, Dulbecco’s Modified Eagle Medium (DMEM) with GlutaMAX™ (± HEPES buffer), Leibovitz L-15 medium with GlutaMAX™, Gibco CO_2_ independent medium, RNA*later*™, foetal bovine serum (FBS), horse serum, goat serum, LIVE/DEAD™ viability kit, Alexa Fluor-488 nm Phalloiden, Alexa Fluor-488 nm goat anti-rabbit secondary antibody, DAPI solution (1 mg/mL), ProLong™ gold antifade mountant, SYBR Green universal master mix, PageRuler™ and high-capacity cDNA reverse transcription kit were purchased from ThermoFisher Scientific (Altrincham, UK).

The custom lentiviral overexpression vector for Heat Shock Protein 10 (HSP10-T2A-EGFP) and control vector were designed and packaged by VectorBuilder GmbH. PrimePCR SYBR Green qPCR primers (HSP10 and RSP18) and 4–20 % Mini-PROTEAN® TGX™ Precast Protein Gels were purchased from Bio-Rad (Watford, UK). Rabbit polyclonal antibodies to HSP10 (Ab53106), Green Fluorescent Protein (GFP) (Ab6556) and sarcomeric α-actinin (SαA) (Ab137346) were purchased from Abcam (Cambridge, UK). Rabbit monoclonal antibodies to GAPDH (G9545) were purchased from Sigma Aldrich (Dorset, UK). IRDye® 800CW goat anti-rabbit secondary antibodies were purchased from LI-CORbio (Cambridge, UK).

The flight hardware and associated consumables were designed, manufactured or procured by Kayser Space Ltd (Didcot, UK). Unless stated otherwise, all other chemicals used in this study were obtained from Sigma Aldrich, Dorset, UK.

### Cell Culture

Human immortalised myoblasts (AB1167_CTL) were routinely maintained in a complete growth medium consisting of PromoCell skeletal muscle basal medium (phenol red-free) supplemented with 20 % (v/v) FBS, 50 μg/mL bovine fetuin, 10 ng/mL epidermal growth factor, 1 ng/mL basic fibroblast growth factor, 10 μg/mL recombinant human insulin, 0.4 µg/mL dexamethasone, 10 μg/ mL gentamicin and 4 mM L-glutamine ^19,20^.

Cells were incubated in a humidified environment at 37 °C with 5 % (v/v) CO_2_, medium exchanges were performed every 48 hours. All stock cultures (AB1167_CTL) were maintained at <70 % confluence and used between passages 4-8 to prevent visual decline of spontaneous contractions.

### Stable Overexpression of Heat Shock Protein 10 (HSP10) by Lentiviral Transduction

To generate a cell line that stably overexpressed HSP10, AB1167_CTL myoblasts were seeded sparsely into 6-well plates (1×10^4^ cells/well) in complete growth media and allowed to adhere overnight (37 °C with 5 % (v/v) CO_2_). The lentiviral vector (**Fig. 1**) was constructed with an EGFP reporter gene, human HSP10 (HSPE1) target sequence, T2A linker and blasticidin resistance gene inserted into the Open Reading Frame (ORF) site. Transcription was under the control of a human EF1A promotor. The plasmid will hereon be referred to as HSP10-T2A-EGFP.

**Figure 1.**
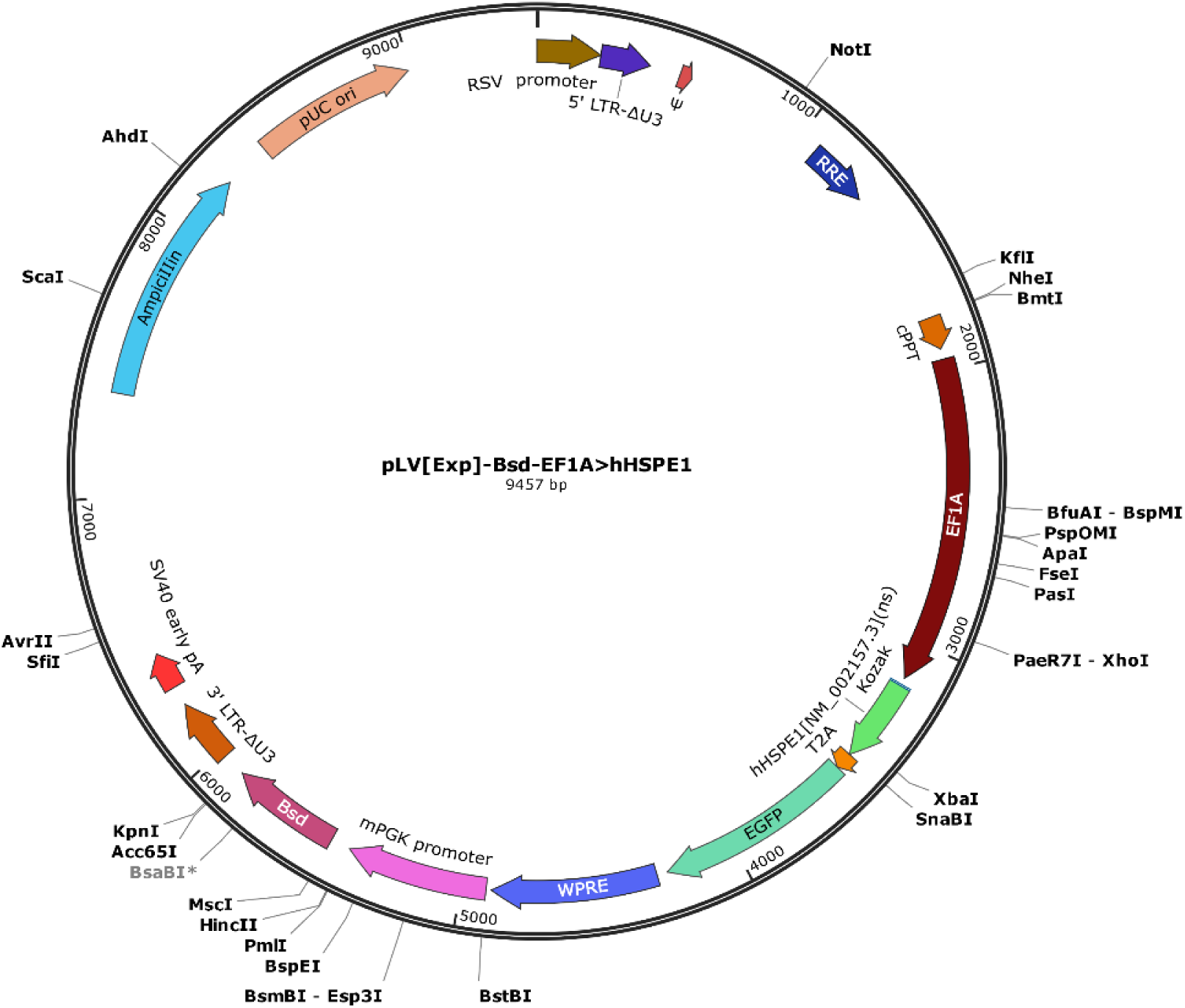
Lentiviral plasmid expression map for the overexpression of HSP10. The plasmid (HSP10-T2A-Bsd) was constructed with a T2A linker to achieve polycistronic expression of an EGFP reporter gene and the human HSP10 (HSPE1) target sequence.

Transgene delivery of HSP10-T2A-EGFP was achieved by lentiviral transduction over a period of 24 hours at a multiplicity of infection (MOI) of 10, 50 and 100, supplemented with 5 µg/mL polybrene. Integration of the viral vector into the host cell genome, ensures consistent transgene expression across cell divisions. The following day, the viral solution was replaced with fresh growth medium, and the cells were incubated overnight before undergoing blasticidin (10 µg/mL) selection for 10 days.

All stock cultures (AB1167_HSP10) were maintained at <70 % confluence and used between passages 4-8 to prevent visual decline of spontaneous contractions. All cultures used for the flight and ground control experiments had >80 % transduction efficiency as confirmed by quantifying the proportion of HSP10-T2A-EGFP expressing cells using a Zeiss Axio Observer Apotome microscope (excitation/emission wavelengths: 488/513 nm).

### Western Blot Analysis of HSP10 Overexpression

In brief, 20 µg of AB1167_CTL and AB1167_HSP10 cell lysates in 1× RIPA buffer underwent electrophoretic separation under reducing conditions using a 20 % Mini-PROTEAN® TGX™ protein gel. The protein was transferred onto a nitrocellulose membrane using a semi-dry transfer method before staining with ponceau red to ensure transfer uniformity. Membranes were blocked in 5 % (w/v) non-fat milk in 1X Tris-buffered saline with 0.01 % (v/v) Tween 20 (TBS-T) for 1 hour at room temperature. Primary antibodies were diluted in 5 % (w/v) non-fat milk 1X TBS-T and incubated with membranes overnight at 4 °C (HSP10 1:500, GFP 1:2500 and GAPDH 1:5000).

Following washes in 1X TBS-T, membranes were incubated with IRDye® 800 C W goat anti-rabbit IgG secondary antibody (LicorBio), diluted 1:10,000 in 5 % non-fat milk TBS-T for 1 hour at room temperature. Membranes were imaged using the Odyssey CLx Imaging System (LicorBio), densitometric analysis was performed using Image J.

### RT-qPCR Analysis of HSP10 Overexpression

RNA was extracted from AB1167_CTL and AB1167_HSP10 cell lysates using a Qiagen RNeasy Plus Kit, following the manufacturer’s protocol. The quality and quantity of the RNA was assessed using a Nanodrop spectrophotometer. Complementary DNA (cDNA) synthesis was conducted using a high-capacity cDNA reverse transcription kit (Applied Biosystems, ThermoFisher).

Amplification of the housekeeping gene and gene of interest was conducted using the ROCHE LC96 light cycler, using a SYBR Green detection system. Primer sequences were proprietary to Bio-Rad and not disclosed, product references: PrimePCR™ SYBR® Green Assay: HSPE1, Human and PrimePCR™ SYBR® Green Assay: RPS18, Human.

### Biofabrication of Human Skeletal Muscle Constructs

Bespoke, biocompatible scaffolds were produced by fused filament fabrication (FFF) 3D printing on an UltiMaker 2^+^ instrument with polylactic acid (PLA) filament. The design solution included internal channels that integrated with the fluid handling circuit of the flight hardware, in addition to three individual reservoirs, each with two anchorage points, around which the muscle constructs spontaneously assemble (**Fig. 2** & **3A**). Post-print, Sylgard 184 (polydimethylsiloxane) blocks were placed at either end of the individual scaffold reservoirs to cover the openings to the internal fluidic channels. The scaffold assemblies were then sterilised in 70 % ETOH before coating in sterile 0.2 % Pluronics**™** F127 overnight (+ 4 °C).

**Figure 2.**
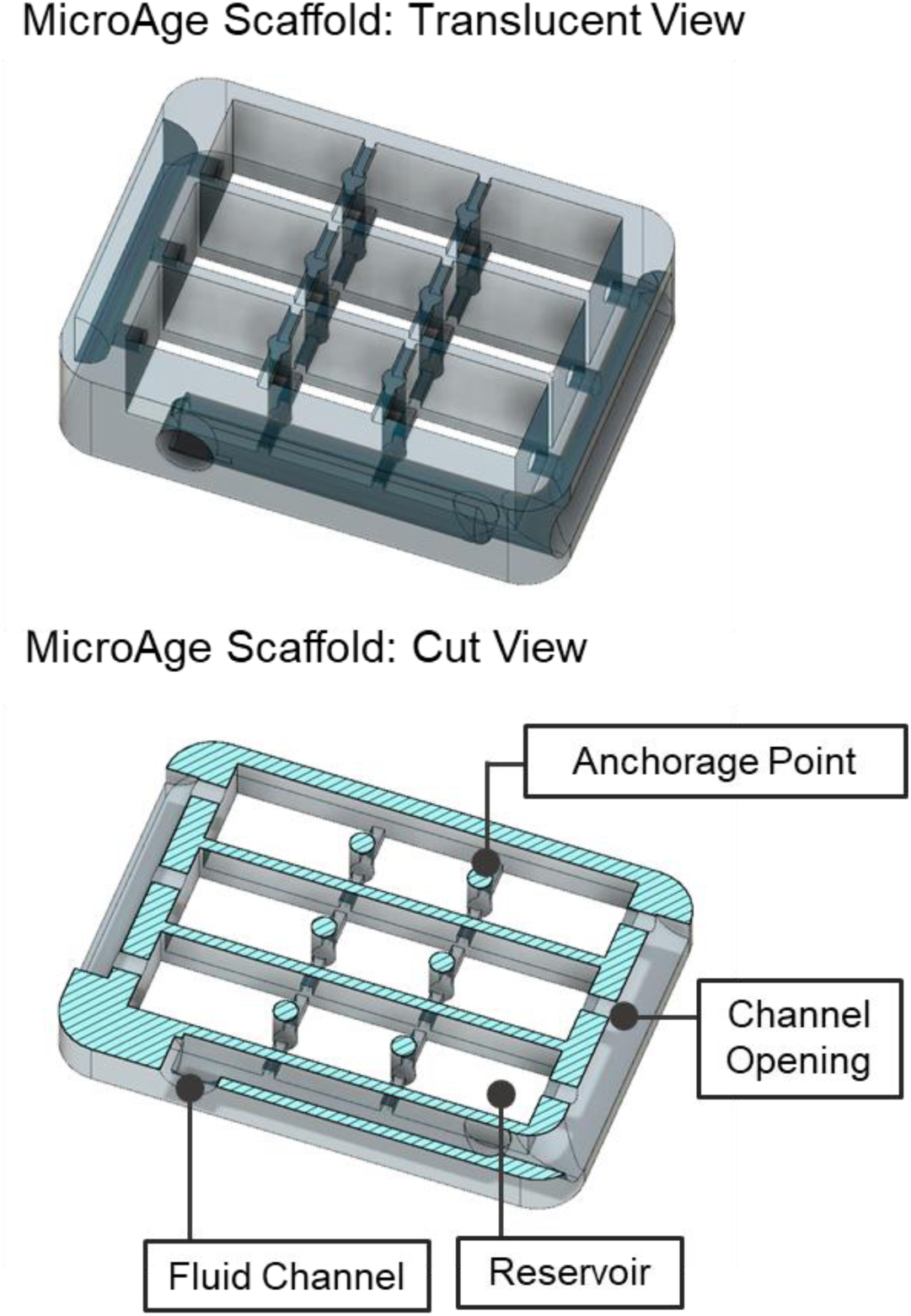
Computer Aided Design (CAD) renderings of the MicroAge scaffold design, highlighting the internal channel architecture, cell seeding reservoirs and anchorage points.

**Figure 3.**
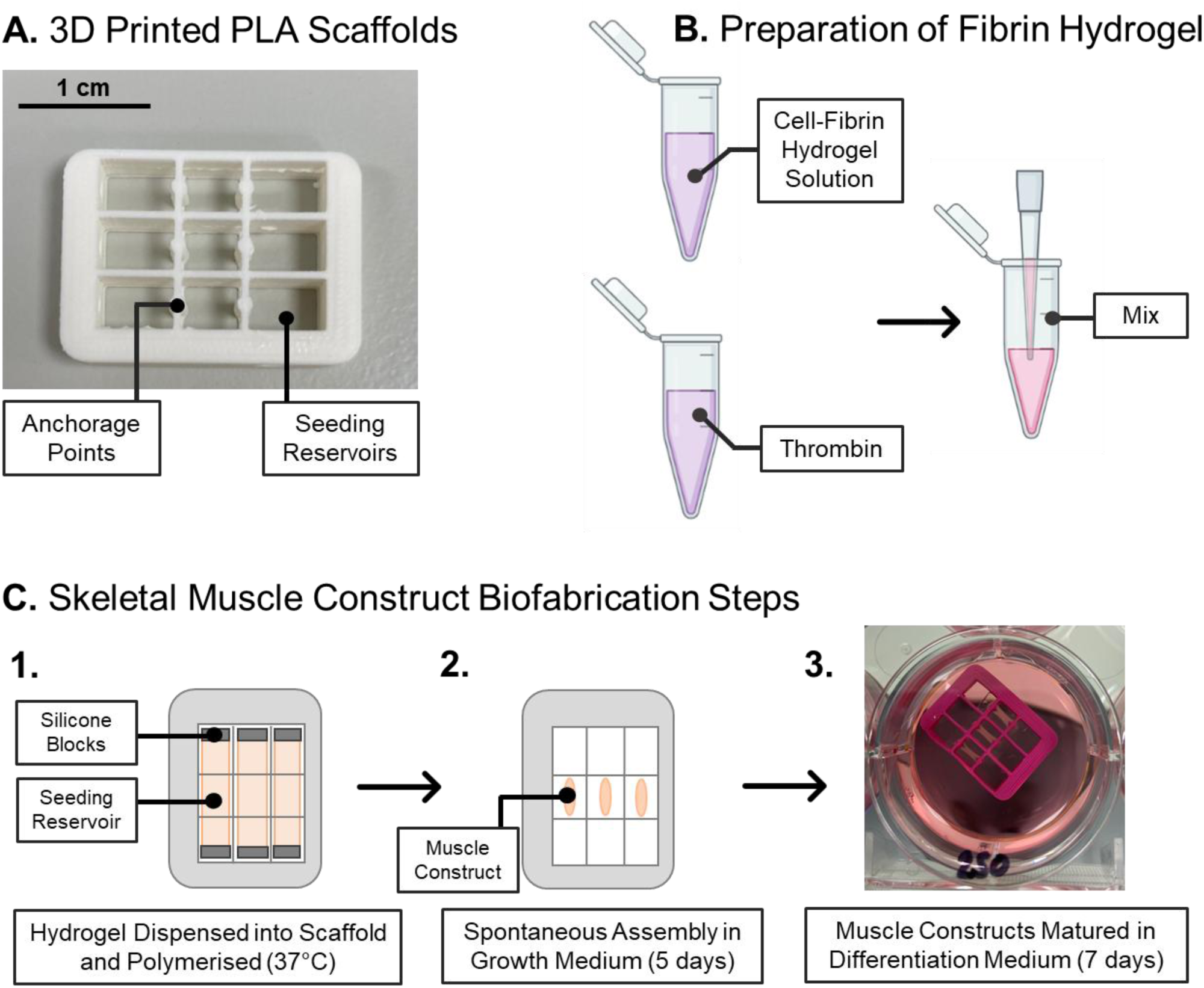
Schematic diagram of the muscle construct biofabrication process. (A) Biocompatible scaffolds were generated by fused filament fabrication on an UltiMaker 2+ instrument. (B) AB1167_CTL or AB1167_HSP10 myoblasts were dissociated by trypsinisation to produce a single-cell suspension before being mixed with a fibrin-hydrogel solution. (C) The hydrogel solution was dispensed into the scaffolds and polymerized at 37 °C for 20 minutes. The muscle constructs were allowed to spontaneously assemble around the scaffold anchor points over 5 days, after which they underwent maturation for a further 7 days.

Expanded myogenic cell populations (AB1167_CTL and AB1167_HSP10), were maintained in complete growth medium, dissociated in 1X trypsin-EDTA to a single cell suspension and encapsulated in a fibrin hydrogel solution before dispensing into the individual scaffold reservoirs at a density of 2.5×10^5^ cells/construct. Each individual scaffold accommodated three muscle constructs.

To prepare one scaffold, a cell solution (7.5×10^5^ cells in 272 μL growth media supplemented with 1.5 mg/mL aminocaproic acid (ACA)) was combined with a hydrogel solution (99 μL growth factor reduced extracellular matrix gel (20 % v/v) and 87 μL fibrinogen (20 mg/mL) in a 1.5 mL polypropylene tube on ice.

To this, 37 μL bovine thrombin was added before pipetting vigorously to mix and dispensing 150 μL of the cell-hydrogel mixture into each of the three scaffold reservoirs (**Fig. 3C**). The cell-hydrogel mixture was polymerised at 37 °C (5 % v/v CO_2_) for 20 minutes followed by incubation in complete growth media supplemented with 1.5 mg/mL aminocaproic acid for five days. During this time, the muscle constructs spontaneously assembled around the static anchorage points within the scaffolds (**Fig. 3C**).

Subsequently, the media was replaced with differentiation medium (DMEM with GlutaMax™ supplemented with 2 % (v/v) horse serum, 10 µg/mL recombinant human insulin, 2 mg/mL ACA and 10 µg/mL gentamicin), for a further 7 days to promote differentiation of myoblasts to myotubes. Routine exchanges of differentiation medium were performed every 48 hours.

### Fluorescent Staining and Imaging of Whole Muscle Constructs

Whole muscle constructs were rinsed with 1X PBS and fixed in 4 % paraformaldehyde (PFA) overnight at 4 °C. Brightfield images were captured on a Zeiss Axio Observer Apotome using the 2.5X objective lens. Resultant images were tiled using Image J.

The muscle constructs were then permeabilized in a buffer containing 0.2 % (v/v) Tween-20 and 0.5 % (v/v) Triton X-100 in 1X PBS for 1 hour at room temperature. Following permeabilisation, constructs were incubated with AlexaFluor 488-conjugated phalloidin (1:20) and DAPI (1:1000), both diluted in the same buffer. Fluorescent imaging was performed using a Zeiss Axio Observer Apotome microscope with a 10X objective lens, using excitation/emission wavelengths of 488/496 nm (AlexaFluor 488) and 350/465 nm (DAPI).

### Cryosectioning and Fluorescent Staining of Muscle Constructs

Muscle constructs were rinsed with 1X PBS and fixed in paraformaldehyde (PFA) overnight at 4 °C, followed by sequential incubation in 10 % and 30 % (w/v) sucrose solutions for 24 hours each. Constructs were then embedded in optimal cutting temperature (OCT) compound and snap-frozen in liquid nitrogen-cooled isopentane.

Longitudinal cryosections (10 µm thick) were prepared using a Leica Biosystems CM1950 cryostat. Sections were rehydrated in 1X PBS and subsequently blocked with a permeabilisation solution containing 0.5 % (v/v) Triton X-100 and 10 % (v/v) goat serum for 1 hour at room temperature. Primary antibodies were diluted in 5 % (v/v) goat serum in 1X PBS and applied to the sections for 1 hour at 4 °C (sarcomeric α-actinin, 1:500). After washing with 1X PBS, sections were incubated with AlexaFluor 488-conjugated goat anti-rabbit secondary antibody (1:500) and DAPI (1:1000), diluted using 5% (v/v) goat serum in 1X PBS.

Sections were mounted using ProLong Gold™ antifade solution before images were captured on a Zeiss LSM 800 confocal microscope with a 40X objective lens, using excitation/emission wavelengths of 488/496 nm (AlexaFluor 488) and 350/465 nm (DAPI).

### Flight Hardware Design Overview

The flight hardware (FH) configuration was composed of two major components; (1) an Experiment Unit (EU) including a Culture Chamber (CC) to house the muscle constructs and (2) the experiment container (EC), based on a standard Kubik Interfacing Container (KIC) footprint, providing an electrical and mechanical interface with the Kubik incubator on board the ISS. The two components together provided two layers of liquid containment (LoC) for the experiment and are described in more depth in the following sections. The materials used to manufacture the EU were selected based on their widely known biological compatibility and excellent chemical resistance.

### Experiment Unit Design

The MicroAge EU can be broken down into four core subsystems, the CC, reservoirs, electronics and the peristaltic pump assembly (**Fig. 4A**), a detailed summary of critical EU components is provided in **Table 1**. The muscle constructs were housed within the CC, anchored to their 3D printed scaffolds. A gas permeable membrane and vented CC lid permitted gas exchange with both the free air volume of the EC and Kubik, providing an aerobic environment to support cellular respiration (**Fig. 4B**). The CC was sealed by securing the gas-permeable membrane between a silicone gasket and a stainless-steel lid. The lid featured vent holes and was fixed in place using counter-sunk stainless-steel screws, creating one LoC.

**Figure 4.**
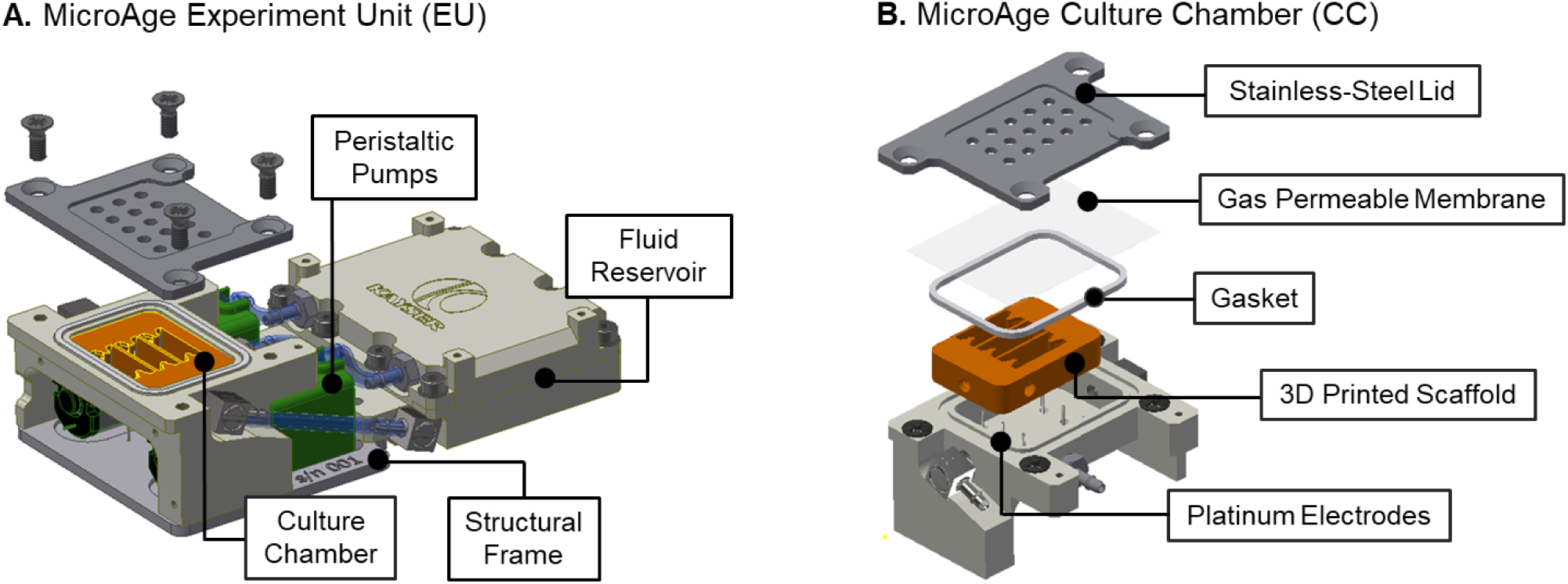
CAD renderings displaying the core components of the Experiment Unit (EU) Design. (A) The EU is composed of the Culture Chamber (CC), fluid reservoir, peristaltic pump assembly and electronics. (B) The CC accommodates the muscle constructs anchored to 3D printed scaffolds. The CC can be further subdivided into its constituent parts including a stainless-steel lid, gas permeable membrane, silicone gasket, platinum electrodes and stainless-steel barbs. The whole EU assembly fits within the Experiment Container (EC).

**Table 1.**
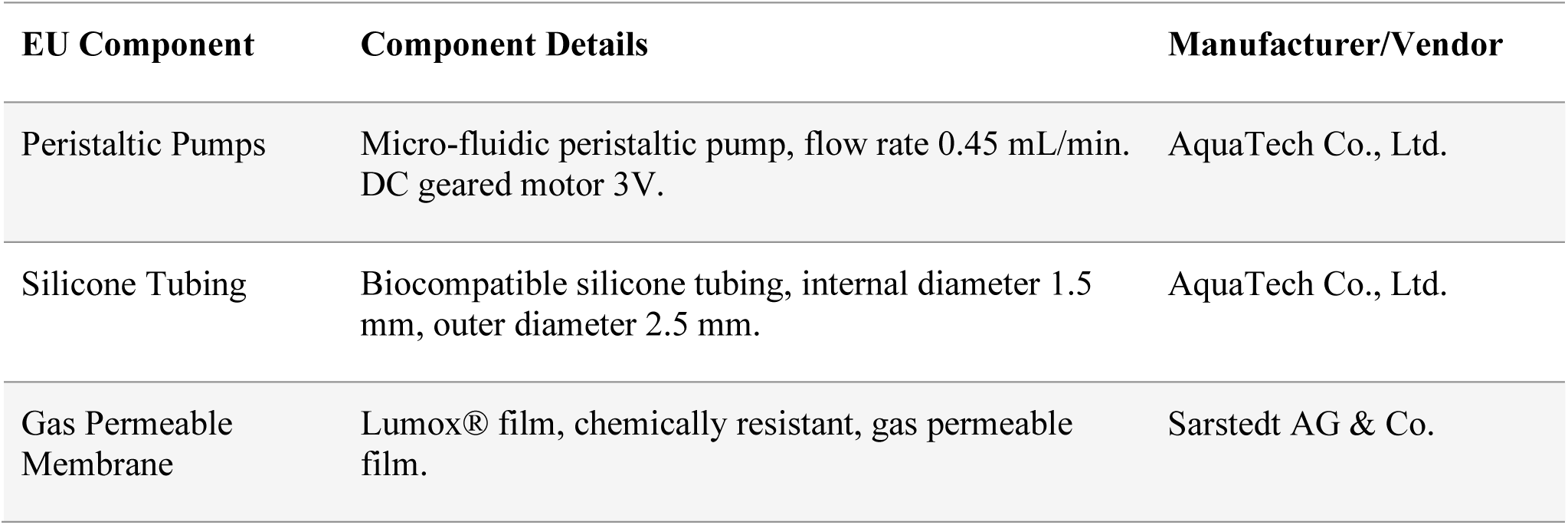
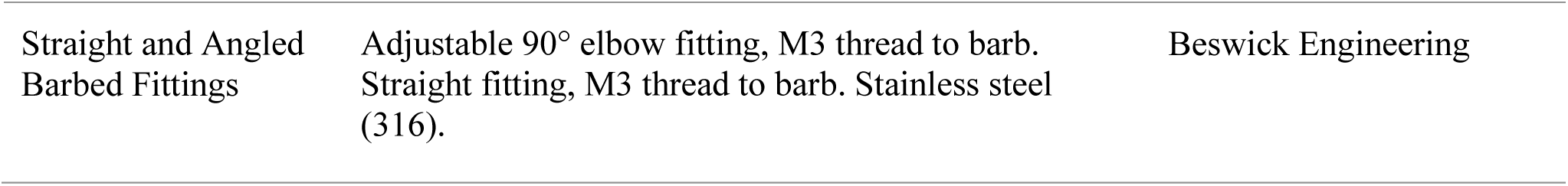
Details of critical EU components.

| **EU Component** | **Component Details** | **Manufacturer/Vendor** |
| --- | --- | --- |
| Peristaltic Pumps | Micro-fluidic peristaltic pump, flow rate 0.45 mL/min. DC geared motor 3V. | AquaTech Co., Ltd. |
| Silicone Tubing | Biocompatible silicone tubing, internal diameter 1.5 mm, outer diameter 2.5 mm. | AquaTech Co., Ltd. |
| Gas Permeable Membrane | Lumox® film, chemically resistant, gas permeable film. | Sarstedt AG & Co. |
| Straight and Angled Barbed Fittings | Adjustable 90° elbow fitting, M3 thread to barb. Straight fitting, M3 thread to barb. Stainless steel (316). | Beswick Engineering |

The CC body was made from machined PEEK (polyether ether ketone), with fluid handling channels, epoxy cemented platinum electrodes (99.99 % purity, 0.5 mm diameter) and mounting points for the pump structural frame. All components of the CC that were in contact with the samples or culture fluids were biocompatible and sterilised by autoclave. During the flight experiment, each muscle construct was electrically stimulated by two platinum electrode pins (six pins per CC) which were attached to a printed circuit board (PCB) underneath the CC. The electrodes were arranged in the CC such that they did not interfere with the sample but were close enough proximity (∼1 mm) to electrically stimulate each construct to contract.

The fluid reservoir was a two well design, allowing for the storage of two different liquids when sealed (culture medium and fixative). The reservoir was sufficiently sized so that two medium refreshes and one fixative (PBS or RNALater™) flush could be performed per EU. Inside the reservoir was a deformable membrane, which served two purposes. Firstly, it allowed for fluids to be pumped in microgravity, acting as a bladder it forced the fluid towards the outlet. Secondly it could deform, allowing the waste medium from the CC to be stored in the reverse side of the same well, separated from the fresh fluid. The membrane was designed with a gasket, therefore guaranteeing one LoC.

The EU pump assembly was composed of the peristaltic pumps, tubes, connectors and structural frame, connecting the CC to the reservoir via silicone tubing and stainless-steel barb fittings. Peristaltic pumps were chosen as the optimal system for moving fluids as the culture fluids only come into contact with the inside of the sterile silicone tubing, they could deliver a flow rate of 0.45 mL/minute, and the pumps acted as non-return valves when unpowered, reducing the risk of cross-contamination between the fresh and spent media. The pumps slotted into an aluminium structural frame and were secured to the bottom of the CC using counter sunk stainless-steel screws.

### Electrical Stimulation and Impedance Sensing System

The primary functions of the EU electronics board were four-fold: (a) experiment timeline execution, (b) electrical stimulation of the muscle constructs to contract following a specific contraction protocol, (c) detection of muscle contractions by means of electrochemical impedance spectroscopy and (d) data storage.

The programmed electrical stimulation sequence occurred over a 15-minute period and consisted of 180 segments (clusters), lasting 5 seconds each (thus total time of 15 minutes), delivered by the six platinum electrodes embedded in the base of each CC. Each segment delivered 500 ms of electrical stimulation and a 4.5s period without stimulation. Within the stimulation period (500 ms), the muscle constructs were exposed to 25 bipolar square wave pulses (1 ms +10 V → 1 ms -10 V) at 50 Hz. This results in one single muscle contraction that lasts ∼700 ms (i.e. 500ms of contraction during the electrical stimulation and ∼200 ms allowed for relaxation of the construct).

In the time between stimulation pulses, a low intensity ∼200 mV AC signal was used to measure the impedance between the electrodes. One frequency was measured between each stimulation pulse, giving a total 25 impedance measurements (frequency range 1 KHz → 100 KHz), After each stimulation pulse the system was given 6 ms to stabilise, then 20 consecutive measurements were taken over a period of 2 ms and averaged. The readings were stored in the non-volatile memory. Impedance measurements continued in the 4.5s between contractions cycling through one frequency every 20 ms.

### Experiment Container Design

The EC was comprised of a lid and body which provided a single LoC for the experiment, with the LoC being maintained through the compression of the silicone gasket by eight screws (**Fig. 5**). Anodized, high stress-corrosion and crack resistant aluminium was used to manufacture the EC to protect it against surface corrosion and cosmetic damage. The interface between the EC and Kubik incubator was via its electrical connector, suppling the EU with power to run the experimental sequence autonomously via firmware.

**Figure 5.**
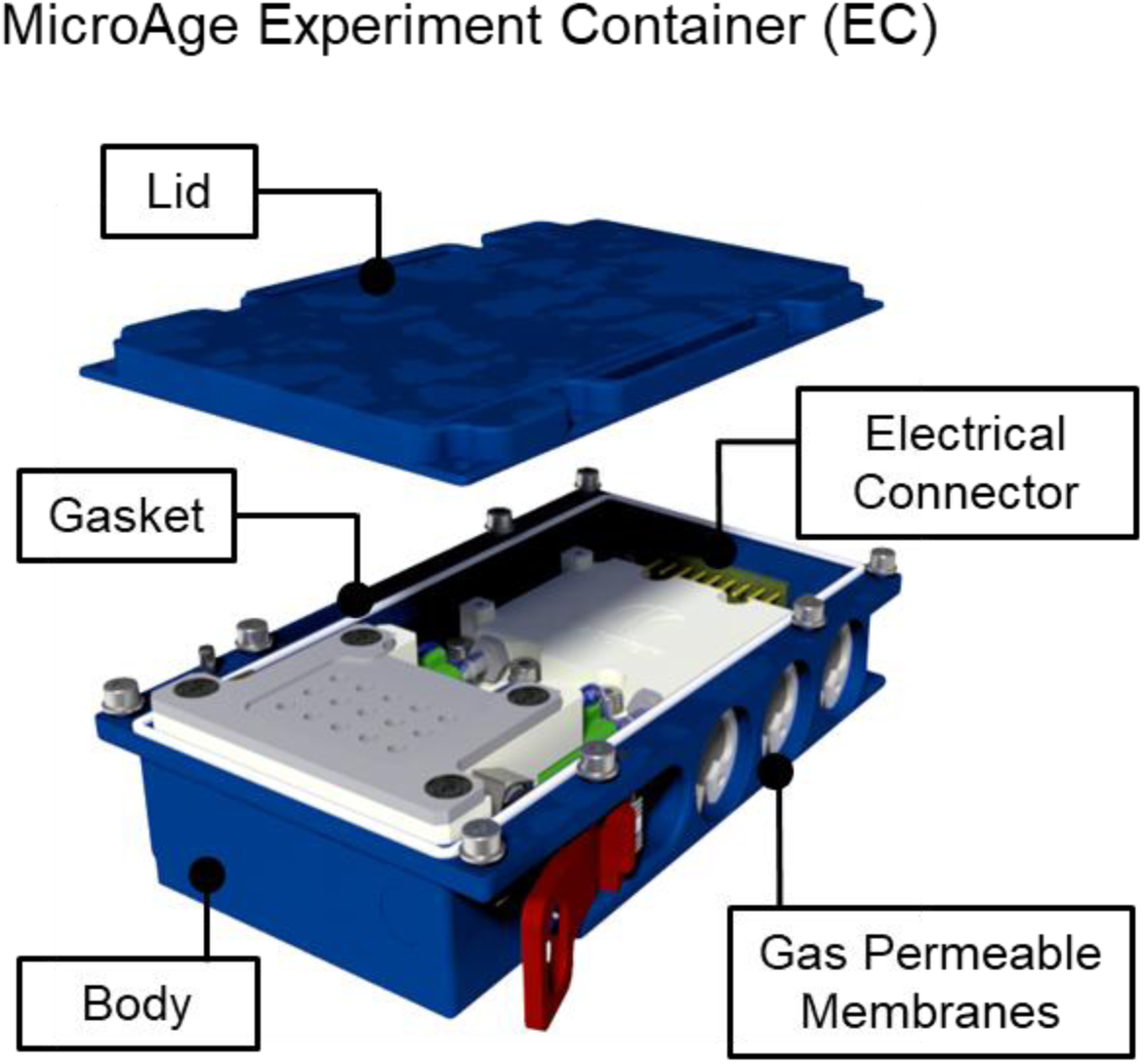
Overview of the MicroAge Experiment Container (EC), highlighting the constituent components of the design solution.

The design of the EC allowed for free gas exchange with Kubik through three gas permeable silicone membranes. This method maintains the LoC as the membrane only permits gas molecules to pass. The membrane was sealed to the EC using an ‘O-ring’ and compressed using an elastic, steel ring retainer. To prevent any external damage to the membrane, the assembly is recessed into the EC side wall with a steel mesh plate covering the membrane (**Fig. 5**).

Temperature data loggers (iButtons – DS1922L) were affixed to the external surface of four ECs in order to monitor the mission temperature profile from handover to sample return.

### Flight Hardware Integration into Kubik

Kubik is a temperature-controlled incubator housed on the Columbus module of the ISS. There are several removable inserts available depending upon individual experimental requirements. The centrifuge insert (CI) was selected for the MicroAge experiment as it accommodated 16 ECs in static positions, thus exposing them to the effects of µg, and 8 ECs in the centrifuge positions. The centrifuge was set at 1g during the flight experiment to simulate earth’s gravity with the gravity vector in the centrifuge perpendicular with the long axis of the muscle constructs. Kubik was pre-warmed to 37 ± 0.5 °C prior to use and maintained within the stated range for the duration of the experiment.

### MicroAge Experimental Design and Hardware Configuration in Flight

As displayed in **Table 2**, the flight hardware was allocated to specific sub-experiments based upon the cell type (AB1167_CTL or AB1167_HSP10) and end-point fixative used in each EU, the gravity level to which they were exposed (i.e. their positional arrangement in Kubik), and the electrical stimulation status of the muscle constructs. Each sub-experiment was allocated two EUs housing three muscle constructs, providing six biological samples per analytical arm of the study.

**Table 2.** MicroAge Experiment Unit (EU) groupings.

| Sub-Experiment | Experiment Unit No | Cell Type | Gravity | Electrical Stimulation | Fixation |
| --- | --- | --- | --- | --- | --- |
| SE1 | 1 | AB1167_CTL | $\mu\text{g}$ | ✓ | RNALater™ |
|  | 2 |  |  |  |  |
| SE2 | 3 | AB1167_CTL | $\mu\text{g}$ | ✓ | 1X PBS |
|  | 4 |  |  |  |  |
| SE3 | 5 | AB1167_HSP10 | $\mu\text{g}$ | ✓ | RNALater™ |
|  | 6 |  |  |  |  |
| SE4 | 7 | AB1167_HSP10 | $\mu\text{g}$ | ✓ | 1X PBS |
|  | 8 |  |  |  |  |
| SE5 | 9 | AB1167_CTL | 1g<sub>s</sub> | ✓ | RNALater™ |
|  | 10 |  |  |  |  |
| SE6 | 11 | AB1167_CTL | 1g<sub>s</sub> | ✓ | 1X PBS |
|  | 12 |  |  |  |  |
| SE7 | 13 | AB1167_CTL | 1g<sub>s</sub> | ✗ | RNALater™ |
|  | 14 |  |  |  |  |
| SE8 | 15 | AB1167_CTL | 1g<sub>s</sub> | ✗ | 1X PBS |
|  | 16 |  |  |  |  |
| SE9 | 17 | AB1167_CTL | $\mu\text{g}$ | ✗ | RNALater™ |
|  | 18 |  |  |  |  |
| SE10 | 19 | AB1167_CTL | $\mu\text{g}$ | ✗ | 1X PBS |
|  | 20 |  |  |  |  |
| SE11 | 21 | AB1167_HSP10 | $\mu\text{g}$ | ✗ | RNALater™ |
|  | 22 |  |  |  |  |
| SE12 | 23 | AB1167_HSP10 | $\mu\text{g}$ | ✗ | 1X PBS |
|  | 24 |  |  |  |  |

*Abbreviations: $\mu\text{g}$ , microgravity; 1g<sub>s</sub>, simulated gravity via centrifugation; CTL: Muscle constructs fabricated using human immortalised muscle cells; AB1167\_HSP10: Muscle constructs fabricated using human immortalised muscle cells overexpressing HSP10.*

Upon return of the samples from the ISS, a ground reference experiment was performed, replicating the upload conditions (as recorded by iButton data loggers affixed to the ECs) and the experiment sequence as it occurred during flight. In this scenario, the number of sub-experiments reduced due to the lack of a simulated 1g_s_ condition on ground. The excess units were used as additional replicates in the AB1167_CTL ± electrical stimulation groups (SE 1-2 and SE 9-10).

### Definition of Sample Upload Conditions

Upload of live, mammalian culture material to the ISS required specific considerations to overcome constraints imposed by the launch and space-flight environments. During the MicroAge mission campaign, access to active temperature control facilities and power were not feasible for the launch and upload phase, as is common for many payloads. As such, culture conditions had to be optimized to ensure that the muscle constructs could survive for a period ≤120 hours using stowage assets with passive thermal control capabilities (Double Cold Bags; DCBs), and without access to power to execute autonomous medium refreshes.

Furthermore, neither the launch stowage assets or Kubik could maintain an environmental CO_2_ concentration of 5 % (v/v) to provide optimal growth conditions and maintain the pH of the media within a physiological range, as is commonly used for cell culture. This therefore required careful selection of an alternative ‘CO_2_-independent medium’ which was compatible with the cells and did not compromise viability compared with the standard DMEM-based differentiation medium formulation.

### Treatment of Myotubes

To screen for suitable media formulations and temperature conditions, myoblasts were harvested via trypsinisation using 1X trypsin-EDTA and seeded as monolayers into appropriate culture vessels. Cells were incubated in complete growth medium for 24 hours to support adherence, followed by a 5-day differentiation period in DMEM with GlutaMax™, supplemented with 2 % (v/v) horse serum, 10 µg/mL recombinant human insulin, and 10 µg/mL gentamicin.

The differentiation medium was subsequently replaced with one of the following, in accordance with the specified exposure time and temperature conditions of the experiment: (1) DMEM with GlutaMAX™ supplemented with 25 mM HEPES, 2 % (v/v) horse serum, 10 µg/mL recombinant human insulin, and 10 µg/mL gentamicin; (2) Leibovitz L-15 medium with GlutaMAX™, supplemented with 10 mM galactose, 2 % (v/v) horse serum, 10 µg/mL recombinant human insulin, and 10 µg/mL gentamicin; or (3) Gibco CO₂-independent medium, supplemented with 4mM L-glutamine, 2 % (v/v) horse serum, 10 µg/mL recombinant human insulin, and 10 µg/mL gentamicin.

### LIVE/DEAD™ Cell Viability Assays

Monolayers were washed with Hank’s Balanced Salt Solution (HBSS) before being incubated with SYTO 10 green-fluorescent nucleic acid stain (live) and red-fluorescent ethidium homodimer-1 stain (dead) for 15 minutes at room temperature. Following incubation, the cells were washed with HBSS before fixation with 4 % (v/v) glutaraldehyde. Cells were visualised using a Nikon Eclipse TE2000 Microscope using excitation and emission wavelengths of 483/503 nm (SYTO 10) and 528/617 nm (ethidium homodimer-1).

### Overview of the In-Flight Experiment Sequence

The muscle constructs were integrated into the experiment units, after which the ECs were sealed and placed into temperature-conditioned double cold bags (DCBs) for handover and upload. Once on board the ISS, the ECs were removed from the DCBs and installed into Kubik. Kubik then supplied power to the 24 experiment units which permitted autonomous execution of the programmed internal experiment sequence, the sequence was staggered due to Kubik power budget limitations. The program consisted of a series of medium refreshes, a period of electrical stimulation and fixative flushes (**Fig. 6**).

**Figure 6.**
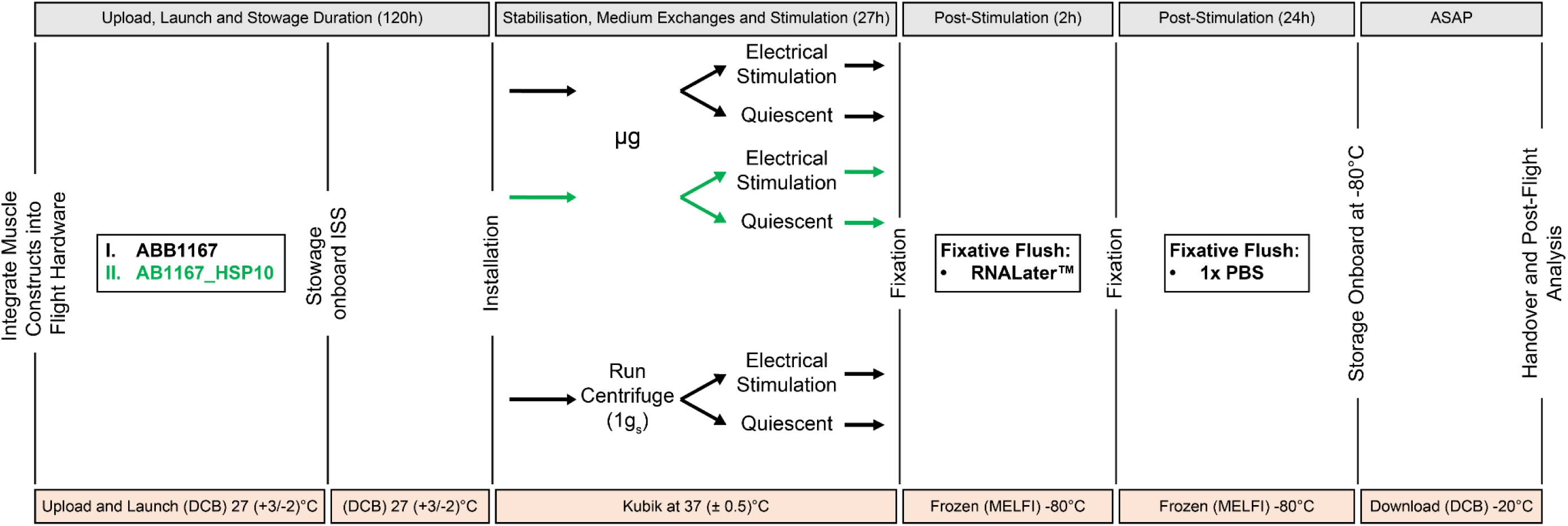
Schematic illustration of the overarching experiment sequence starting with integration of the muscle constructs into the experiment units, upload and launch, installation and experiment execution, fixation and finally, onboard stowage and download. The diagram depicts both the timings (grey bars) and temperature profiles (orange bars) for each step. Abbreviations: DCB, Double Cold Bag; MELFI, Minus Eighty Degree Laboratory Freezer; µg, microgravity; 1gs, simulated 1g (centrifuge); PBS, Phosphate Buffered Saline

## 3 Results

### Bioengineered Skeletal Muscle Constructs

Bioengineered skeletal muscle constructs were generated as previously described in Tollitt *et al.,* 2025, whereby liquid cell-ECM/fibrin hydrogel suspensions were deposited into the reservoirs of 3D-printed PLA scaffolds^21^. Over 12 days, the constructs compacted around internal anchorage points (**Fig. 7A**). To evaluate myotube alignment, the entire muscle constructs were fixed in 4 % (v/v) paraformaldehyde (PFA) and co-stained with Alexa Fluor (488 nm)-conjugated phalloidin and DAPI to visualise the filamentous actin (f-actin) network within individual myotubes.

**Figure 7.**
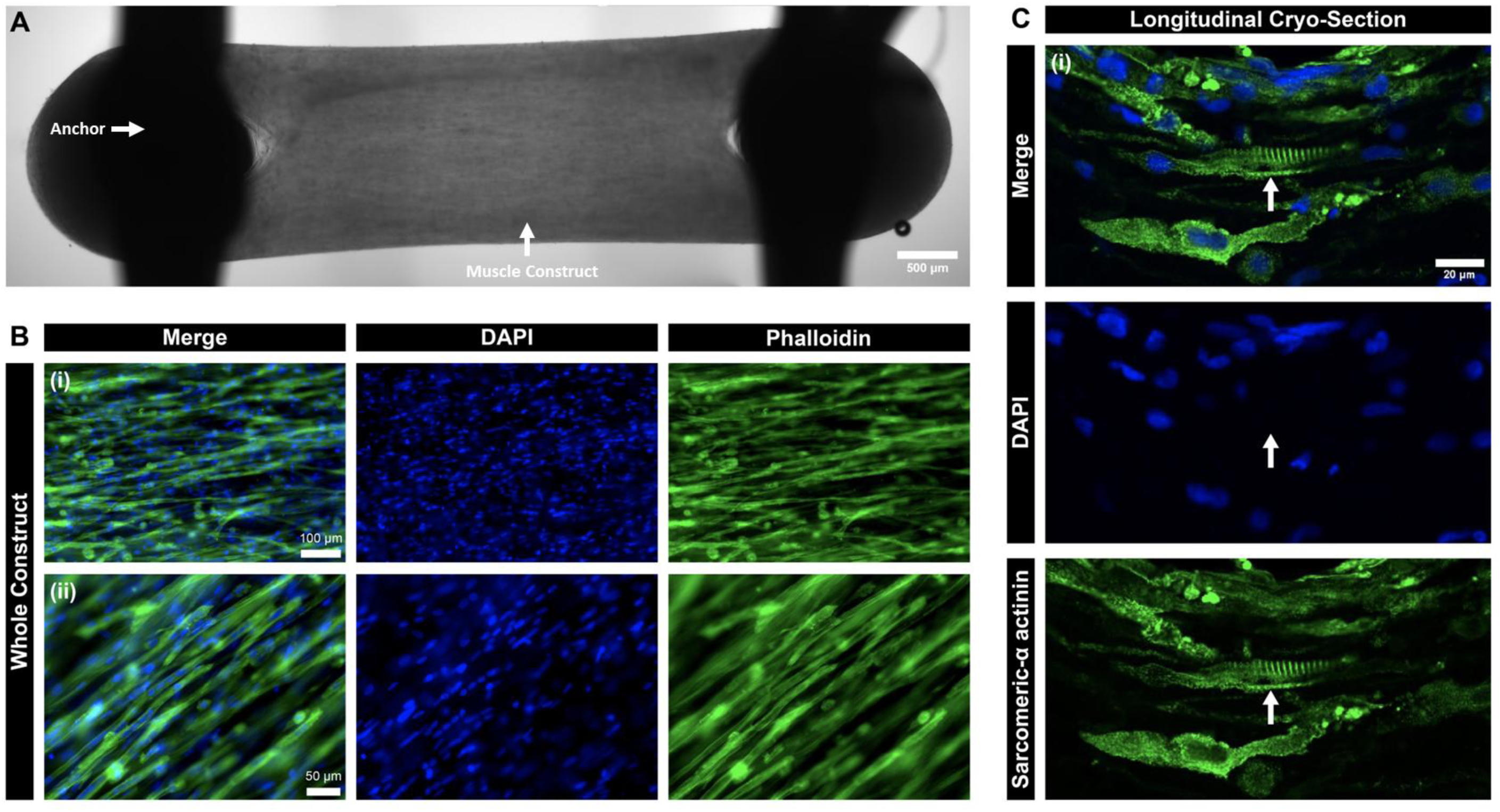
Assessment of skeletal muscle construct architecture. (A) Representative brightfield image of a whole muscle construct fixed in 4 % paraformaldehyde (PFA), scale bar 500 µm. (B) Assessment of myotube alignment across the long axis of whole, fixed (4 % PFA) muscle constructs. Muscle constructs were stained with Alexa Fluor 488-conjugated phalloidin to highlight the filamentous actin (f-actin) networks in individual myotubes and DAPI to highlight the nuclei, scale bar (i) 100 µm and (ii) 50 µm. (C) Longitudinal cryo-sections of muscle constructs, co-stained to detect sarcomeric-α actinin and nuclei (DAPI), scale bar 20 µm, arrow highlights sarcomeric striations.

Differentiated muscle constructs demonstrated end-to-end alignment of multinucleated myotubes along the construct’s longitudinal axis, as depicted in **Figs. 7A-B**. Longitudinal cryosections of mature muscle constructs were stained with sarcomeric α-actinin, a structural protein essential for sarcomere stability and organisation ^22^, revealing the characteristic striated pattern indicative of the organised arrangement of contractile proteins (**Fig. 7C**).

### Stable Overexpression of HSP10 in Human Immortalised Myoblasts

To generate stable overexpression of HSP10 in human skeletal myoblasts, the lentiviral transduction efficiency of the HSP10-T2A-EGFP plasmid was examined at 24, 144 and 240 hours after blasticidin application and at different multiplicities of infection (MOIs) to select for optimum conditions (10-100; **Fig. 8A-C**). Transduction efficiencies were calculated by counting the proportion of cells that expressed HSP10-T2A-EGFP plotted as a percentage of the total number of cells in the field of view (FOV), The absence of auto-fluorescence was confirmed in non-transduced AB1167_CTL myoblasts (data not shown).

**Figure 8.**
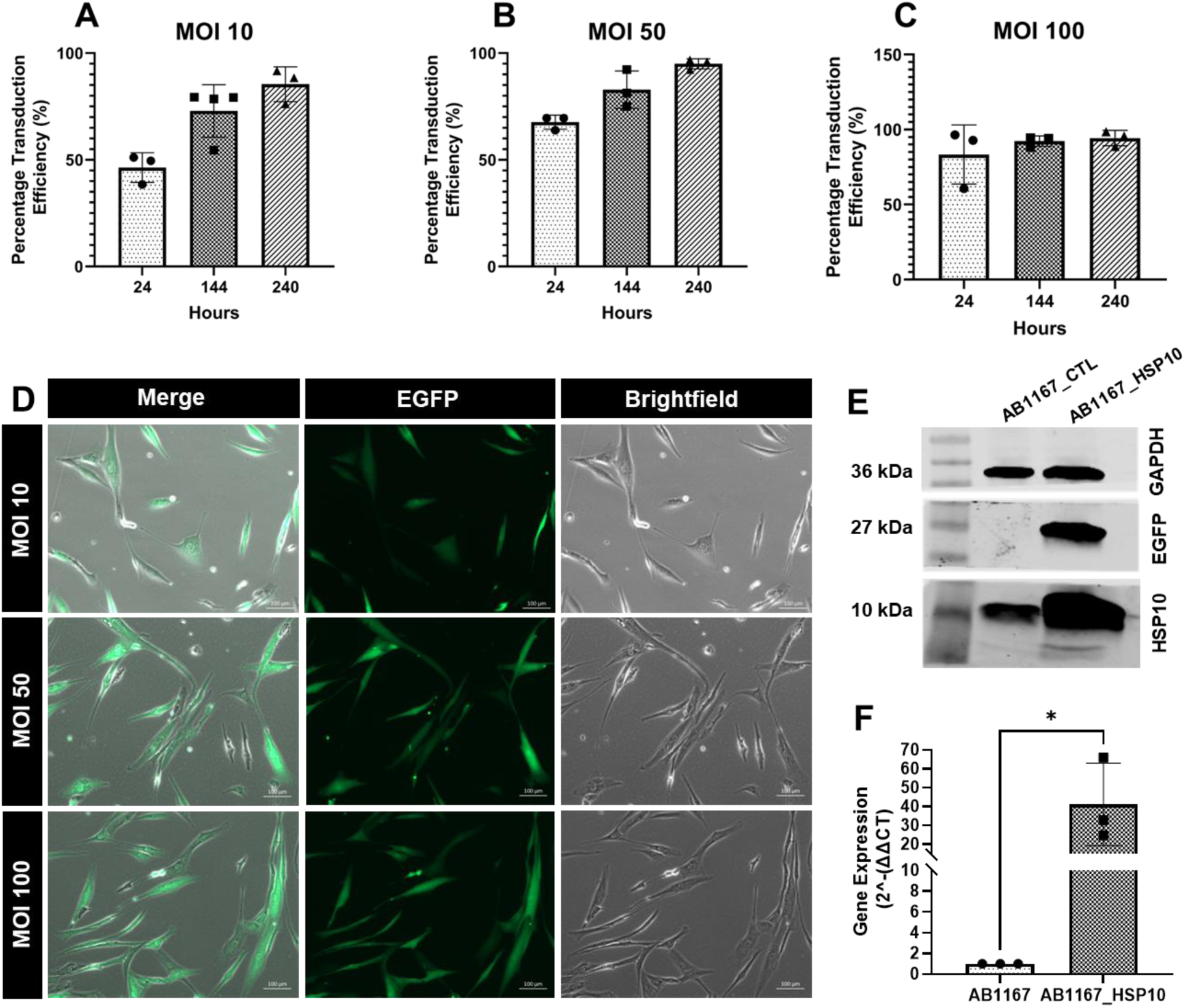
Optimising the stable overexpression of Heat Shock Protein 10 (HSP10) in human immortalised myoblasts (AB1167_CTL) by lentiviral transduction. (A-C) Percentage transduction efficiency was calculated by counting the number of cells that expressed the fluorescent EGFP reporter as proportion of the total number of cells within a field of view (FOV), a minimum of three separate FOVs were analysed per condition. (D) Representative (n of 3) brightfield and fluorescent (excitation 488nm, EGFP reporter) images of the AB1167_HSP10 myoblasts after 240 hours in blasticidin selection media (MOI 10-100), scale bar 100µm. (E) Representative western blot image demonstrating that HSP10 is overexpressed in the mutant cell line, as well showing the absence of EGFP in the control cell line (MOI 100). GAPDH was used as the loading control. (F) Gene expression data (RT-qPCR), illustrating increased expression of the HSP10 gene in the mutant cell line (MOI 100). All graphical values are expressed as a mean ± standard deviation (minimum n=3). Data were tested for Gaussian distribution using a Shapiro-Wilk test before statistical significance was determined using a student’s t-test with Welch’s correction. *p value < 0.05.

At 24 hours, a MOI of 10 had a transduction efficiency of 46.3 ± 6.8 % which increased to 85.4 ± 8.1 % after 240 hours in the blasticidin selection media. A MOI of 50 resulted in a higher efficiency at 24 hours (67.7 ± 3.3 %), increasing to 95 ± 2.3 % after 240 hours under selection. The greatest starting efficiency was yielded from a MOI of 100 (83.3 ± 19.7) which reached 92.4 ± 3.4 % after 144 hours, this effect was also clearly seen following widefield imaging of the EGFP plasmid in cells (**Fig. 8D**). As such, this condition was selected to produce the stably transduced, mutant cell line (AB1167_HSP10).

To further characterise the cell line, whole cell lysates were subject to analysis via western blot (**Fig. 8E**). Results showed the presence of the EGFP reporter in the AB1167_HSP10 cells as well as a greater abundance (212.6 % increase) of the HSP10 protein compared with the AB1167_CTL. Gene expression data further confirmed the upregulation of HSP10 by 41.1 ± 21.8-fold in the AB1167_HSP10 line compared with control (**Fig. 8F**).

### Optimisation of Sample Upload Conditions: CO_2_-Independent Medium Selection

A series of CO₂-independent medium formulations were screened using AB1167_CTL myotubes in two-dimensional culture to assess performance over an 120-hour period (**Fig. 9B-D**). Standard DMEM-based differentiation medium (**Fig. 9A**) was used as a control (37◦C, 5 % (v/v) CO_2_). To simulate passive upload scenario to the ISS, cells were tested under two conditions: a ‘fasted’ state, with no medium exchanges and a ‘fed’ state, where medium exchanges were performed at regular (48 hour) intervals (37◦C, ambient CO_2_). End-point measuresments of cell viability were assessed using a LIVE/DEAD™ cell imaging kit.

**Figure 9.**
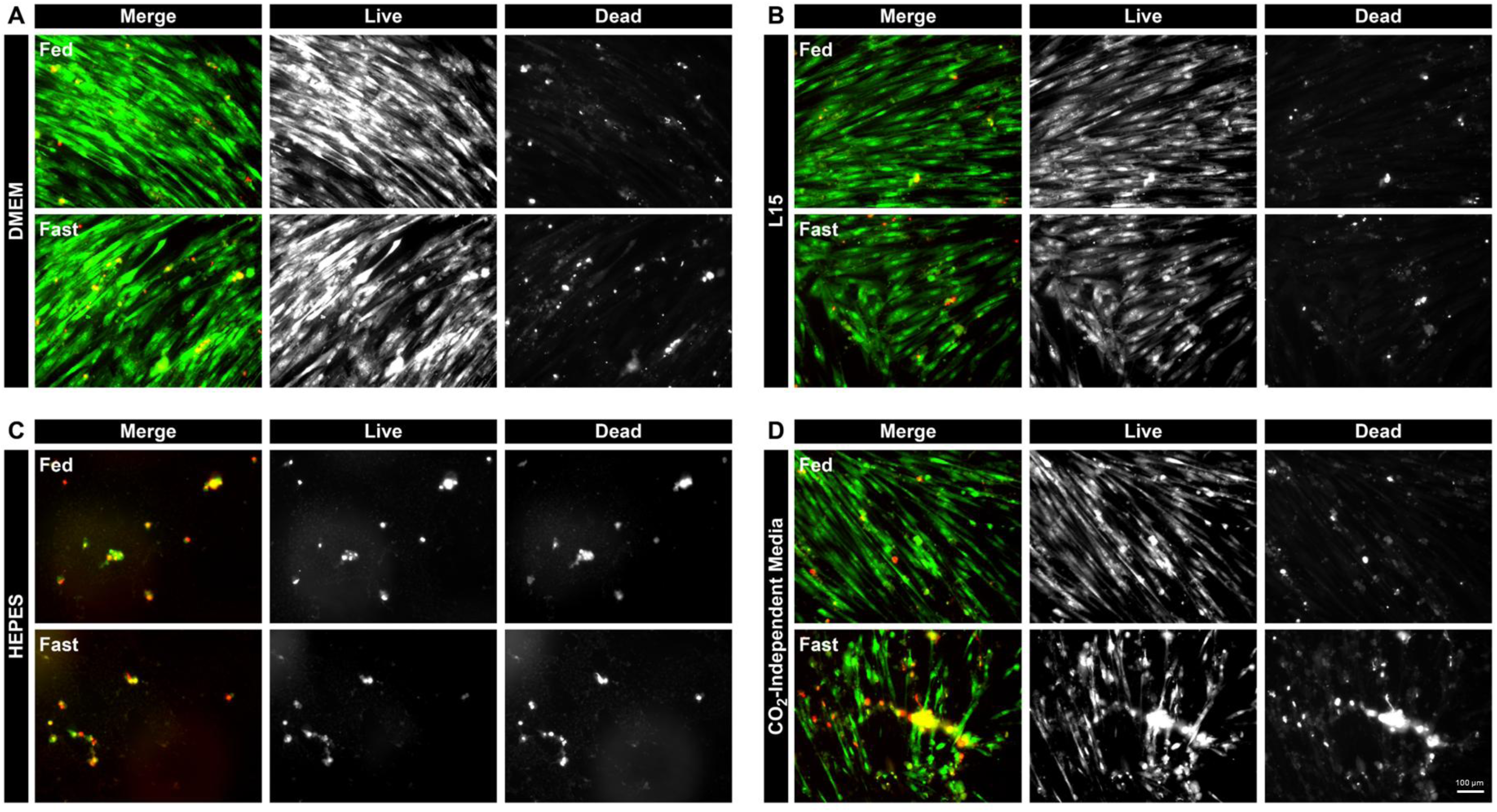
Representative images (n=3) LIVE/DEAD™ assays evaluating the performance of different media formulations, that do not require 5 % (v/v) CO₂ to maintain physiological pH, using AB1167_CTL cells over a 120-hour period (37 °C). Two test conditions were assessed: a ‘fasted’ state without medium exchanges to simulate passive upload conditions and a ‘fed’ state with medium exchanges every 48 hours. (A) DMEM-based differentiation medium under standard culture conditions (37 °C, 5 % CO₂) was used as the control. (B) Leibovitz L-15 medium with GlutaMax™, supplemented with 10 mM galactose, 10 µg/mL recombinant human insulin, and 2 % horse serum. (C) HEPES-buffered DMEM with GlutaMax™, supplemented with 25 mM glucose, 10 µg/mL recombinant human insulin, and 2 % horse serum. (D) Gibco CO₂-independent medium, supplemented with 4 mM L-glutamine, 10 µg/mL recombinant human insulin, and 2 % horse serum.

The DMEM controls performed as expected, with the myotubes in both the ‘fed’ and ‘fasted’ states remaining viable throughout the culture period. Minimal cell death occurred due to the absence of medium exchanges up to 120 hours (**Fig. 9A**). In contrast, myotubes cultured in HEPES-buffered DMEM under ambient CO_2_ conditions exhibited a complete loss of viability, even when medium exchanges were performed at regular intervals (**Fig. 9C**).

Both the Gibco CO_2_-independent medium (**Fig 9D**) and Leibovitz L15 medium (**Fig. 9B**) performed well under ambient CO_2_ levels_,_ particularly when the cells were receiving regular medium exchanges. However, in the absence of medium refreshes, moderate cell death and cell detachment was observed with the Gibco CO_2_-independent medium. As such, the Leibovitz L15 medium was selected for use going forward.

### Optimisation of Sample Upload Conditions: Temperature Control

During upload and prior to insertion into Kubik, the ECs were stowed for launch in double cold bags (DCBs). The DCBs are used routinely for passive thermal control of science payloads and are able to maintain a relatively broad temperature range for at least 120 hours through the use of pre-conditioned thermal bricks. This was to account for potential launch delays and time to ISS docking.

To determine an optimal thermal range for the cells during the upload process, various temperature set points ranging from +4 °C to +37 °C were assessed using Leibovitz L15 medium over the course of 120 hours. During this period, medium exchanges were not performed, and AB1167_CTL myotubes were maintained in two-dimensional culture. For comparison, control myotubes were maintained using a standard DMEM-based differentiation medium at 37 °C (5 % (v/v) CO_2_), with medium exchanges performed every 48 hours. Cell viability was assessed at incremental time points (24, 72 and 120 hours) using a LIVE/DEAD™ assay kit. The results are presented in **Fig. 10A-E**.

**Figure 10.**
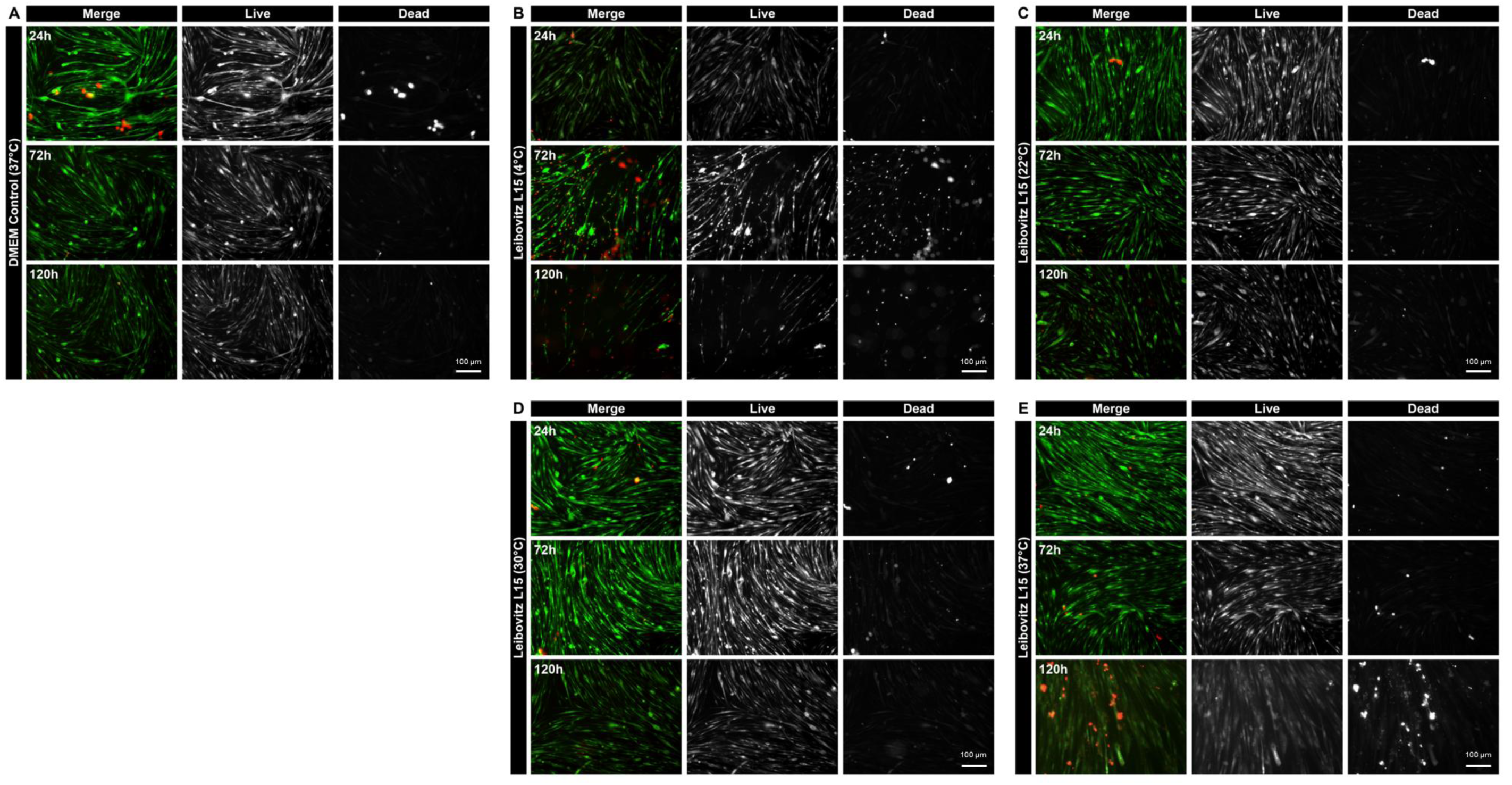
Representative images (n=3) depict temperature tolerance screening of AB1167_CTL myotubes cultured in Leibovitz-L15 medium (ambient CO_2_) at various upload temperature set points (4–37 °C). Cells were incubated over a 120-hour period without medium exchanges (fasted) to mimic worst-case, passive upload conditions to the ISS. LIVE/DEAD™ assays were conducted at 24, 72, and 120-hour intervals to evaluate cell viability. The data were compared to a DMEM control (37 °C, 5 % (v/v) CO_2_) across the same culture period, where the medium was refreshed every 48 hours (A). Incubation in Leibovitz-L15 medium at 4 °C (B), 22 °C (C), 30 °C (D) and 37 °C (E).

The DMEM-based control myotubes, which underwent regular medium exchanges, exhibited minimal signs of cell death or detachment throughout the 120-hour culture period (**Fig. 10A**). In contrast, myotubes cultured in Leibovitz L15 medium at +4 °C (**Fig. 10B**) began showing signs of reduced viability as early as 72 hours, with complete cell detachment observed at 120 hours. More favorable results were observed at +22 °C (**Fig. 10C**) and +30 °C (**Fig. 10D**) with myotubes remaining viable across all time points tested. Notably, myotubes cultured for 120 hours at +30 °C exhibited no signs of sparsity, which was more pronounced at +22 °C. Myotubes cultured in Leibovitz L15 at +37 °C (**Fig. 10E**), performed as anticipated across the earlier timepoints, however after 120 hours some cell death was evident.

### Validation of the Electrical Stimulation and Impedance Sensing System

The impedance monitoring system continuously measured the impedance spectrum of the CC during the stimulation phase (**Fig. 11**). Contractions were monitored by calculating the ratio between the impedance measurements during the 0.5s stimulation phase and the impedance measurements at ‘rest’ (4s after stimulation). Samples unaffected by contractions show a flat impedance ratio between the stimulated and rest phases, however, electrical noise from the stimulation signal was distorting the low-frequency response of the system, resulting in a ratio <1 in the region below 10 kHz even without tissue constructs present.

**Figure 11.**
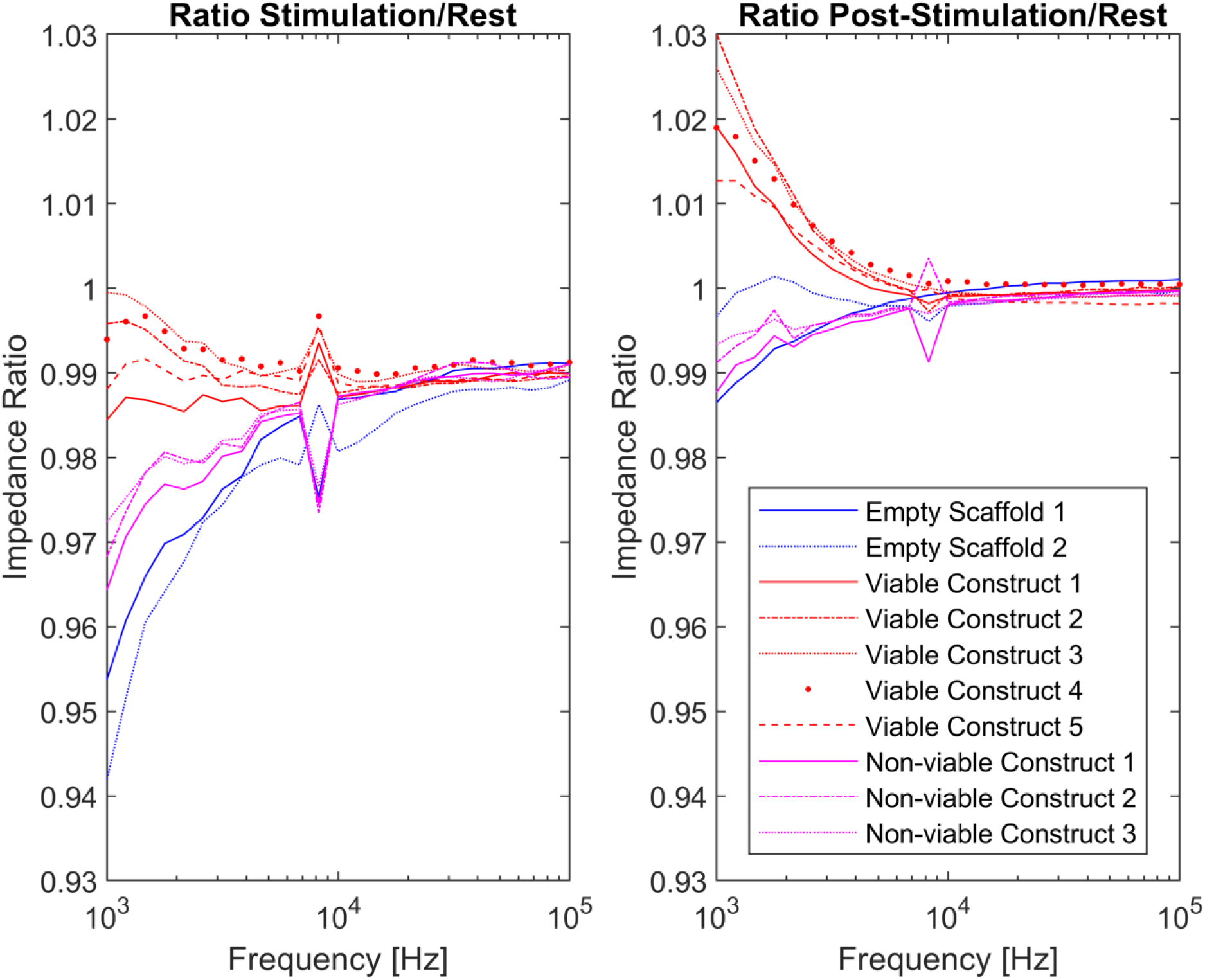
Measurement of the impedance spectrum of the culture chamber containing either empty scaffolds, viable or non-viable AB1167_CTL muscle constructs during the stimulation phase (left) and the post-stimulation phase (right).

As a secondary measure, the ratio between the 0.5s just after stimulation (‘post stimulation’) and the ‘rest’ phase was assessed. Since the impedance changes due to contraction lasts for at least part of the rest period, the signal is still affected by the tissue relaxation but there is much less electrical interference as the stimulation signal is not present. The ratio between this ‘post-simulation’ region and the rested construct stays close to 1 for all frequencies with empty scaffolds with or without non-viable constructs. In all cases, the response of contractile viable constructs deviated from empty scaffolds or non-viable constructs in the region below 10 kHz that allows for reliable detection of whether contractile activity has occurred. A video of the muscle constructs contracting during the electrical stimulation protocol is provided in **Supplementary Fig. 1**.

## 4 Discussion

This publication details the extensive design process and preliminary experiments necessary to create hardware and protocols for maintaining the viability and functionality of human skeletal muscle cells aboard the ISS. This work enabled the study of 3D skeletal muscle constructs, derived from human cells, to assess their responses to microgravity exposure and evaluate the influence of overexpressing the mitochondrial chaperone, HSP10. Furthermore, the developed system facilitated investigations into whether muscle constructs on the ISS responded differently to electrically stimulated contractions compared to ground-based reference experiments, and whether inducing artificial gravity (1g_s_) on the ISS via centrifugation altered microgravity-related responses. Several aspects of this development are notably novel.

### Suitability of Tissue-Engineered Muscle to Study Skeletal Muscle Responses During Space Flight

The pronounced decline in skeletal muscle mass and function seen in humans and animals under microgravity conditions has established skeletal muscle as a primary area of study in this field ^23^. These effects are often studied using orbital platforms like the ISS or ground-based microgravity simulation models such as clinostats and Random Positioning Machines (RPMs) ^24,25^. Large-scale experiments conducted aboard the ISS have encompassed studies of both human and animal muscle, both prior to and following spaceflight, upon return to Earth ^26–28^. Further, there have been numerous studies using cultured muscle cells in various different model systems ^29,30^.

Traditionally these systems have used two-dimensional cultures of mammalian skeletal muscle, derived from satellite cells and driven to differentiate from myoblasts into myotubes adhered to flat, tissue-culture treated surfaces ^31^. A great deal of information has been obtained from such models but the resultant monolayers of myotubes are relatively immature and lack the complex alignment and spatial organisation that reflects native muscle tissue; a limitation that has been increasingly recognised as the field has advanced.

Over the past decade, the biofabrication of 3D skeletal muscle tissue *in vitro* has garnered significant attention. This advancement has enabled both the cultivation and differentiation of human myoblasts into myotubes, encapsulated within synthetic scaffolds or hydrogels engineered to replicate some of the complexities of muscle extracellular matrix (ECM) ^21,32,33^. A comprehensive review of recent trends in skeletal muscle tissue-engineering has been published by Schätzlein and Blaeser, 2022 ^34^.

For the MicroAge Mission, a cell-hydrogel encapsulation method was selected to develop a tissue-engineered human skeletal muscle model that could be anchored onto bespoke 3D-printed, PLA scaffolds^21^. The scaffolds were designed to hold the three muscle constructs in a fixed position, ensuring that the distance between each construct and the platinum electrodes remained consistent across all CCs. This design was crucial for the reliable and reproducible delivery of the electrical stimulation pulses and detection of the resultant impedance signals. Further, the scaffold design featured microfluidic channels that integrated with the flight hardware fluid handling system, facilitating efficient fluid exchanges (≥ 90 %, data not shown) in the CCs (**Figs. 2-3**).

An immortalised human skeletal muscle cell line (AB1167_CTL) was chosen to generate the muscle constructs for this study. This cell line was selected due to its extended proliferative lifespan relative to primary human myoblasts, ability to retain fusion characteristics from the parental cell population and its capacity to express a myogenic program ^19,20^. Additionally, the differentiated myotubes derived from this line have been shown to maintain a functional phenotype, capable of spontaneous contractions as well as contractions following electrical stimulation. This makes the cell line an ideal choice for investigating adaptive responses to contractile activity, particularly in the context of space flight ^21,35^.

As illustrated in **Fig. 7**, the muscle constructs generated from the AB1167_CTL cell line exhibited highly organised, end-to-end alignment of the myotubes along the longitudinal axis of the construct, enhancing the architectural complexity of the model in comparison to myotubes grown in 2D culture. For the MicroAge mission, a naturally derived ECM-based hydrogel formulation was chosen to enhance cell adhesion and growth. This selection was driven by the presence of naturally occurring bioactive motifs, such as arginine-glycine-aspartic acid (RGD), which play a crucial role in promoting cellular interactions and supporting tissue development ^36^. Furthermore, the incorporation of fibrin into the hydrogel formulation introduced binding sites for growth factors, such as basic fibroblast growth factor-2 (bFGF-2) ^37^.

A notable limitation associated with the use of ECM-based hydrogels and fibrin arose from their murine and bovine origins, respectively, which complicated proteomic analyses due to sequence homology with human peptide sequences. Moreover, naturally derived ECM-based hydrogels are subject to inherent batch-to-batch variability, presenting challenges in reproducibility. This issue could potentially be circumvented by using synthetic or defined peptide hydrogels ^38^. Nonetheless, significant variability persists within biofabrication methodologies for engineering skeletal muscle constructs. Critical parameters under continued investigation and optimisation include hydrogel composition, cellular alignment, cell density optimisation, and maturation protocols. For a comprehensive review, refer to Volpi *et al*., 2022 ^39^.

### Efficacy of HSP10 Overexpression in Muscle Constructs

Prior to the mission, it was critical to characterise the AB1167_HSP10 cell line and verify the stable overexpression of the transgene following lentiviral transduction. As shown in **Fig 8**, a high transduction efficiency (92.4 ± 3.4 %) was achieved at an MOI of 100 after just 144 hours of blasticidin selection. Overexpression of HSP10 was confirmed through qPCR and western blot analyses. Importantly, the overexpression did not adversely affect myoblast proliferation or the ability of the myoblasts to fuse and form myotubes in either 2D or 3D cultures (data not shown).

The overexpression of HSP10 in muscle constructs was selected as a proof-of-concept genetic intervention study for the mission, based on its ability to reduce the accumulation of oxidatively damaged proteins (protein carbonyls) and preserve both maximal tetanic force generation and muscle cross-sectional area in murine models of age-related skeletal muscle decline ^17^. HSP10 functions predominantly as a mitochondrial co-chaperone, facilitating the proper folding of proteins within the mitochondrial matrix. This role is particularly crucial under conditions of cellular stress, where protein misfolding may be more prevalent ^40^. Its role in maintaining mitochondrial protein quality control is critical for cellular homeostasis, especially in tissues with high metabolic demands, such as skeletal muscle ^41^.

Given the well-documented role of mitochondrial dysfunction in spaceflight-induced tissue dysfunction ^26^, HSP10 overexpression may present therapeutic potential for mitigating muscle atrophy in such environments. The combination of HSP10 overexpression with physical exercise could further enhance these protective effects. The MicroAge mission aimed to investigate the effects of HSP10 overexpression, both in the presence and absence of electrically induced muscle contractions, under microgravity conditions.

### Selection of CO_2_ Independent Medium and Implications for Construct Metabolism

In standard cell culture practices, a humidified CO₂ incubator set to 5 % (v/v) CO₂ is typically used. The CO₂ dissolves into the cell culture medium, where a fraction reacts with H₂O to form carbonic acid. The carbonic acid subsequently interacts with bicarbonate ions present in the medium, stabilising the pH within a physiological range (7.2–7.4) ^42^. As previously discussed, neither the DCB stowage assets for upload or Kubik could maintain an environmental CO_2_ concentration of 5 % (v/v) to provide optimal growth conditions and maintain the pH of the media within a physiological range. Therefore, an alternative CO_2_-independent media buffering system had to be identified and subsequently tested with the cells.

Tests were conducted using three candidate CO₂-independent culture media, with and without regular medium exchanges across a 120-hour period (**Fig. 9**). Among these, two media, Gibco CO₂-independent medium and Leibovitz L15 medium, demonstrated favourable performance when regular medium exchanges were performed. However, in the absence of medium refreshes, the Gibco CO₂-independent medium exhibited cell death and detachment from the plate. As such, the Leibovitz L15 was selected for use during the mission.

Leibovitz L15 medium was originally formulated for use in CO₂-free systems, buffered by a complement of salts, D-galactose, and free-base amino acids, particularly L-arginine, to maintain a physiological pH in culture ^43^. Furthermore, as observed by Chang and Geyer, media supplementation with D-galactose in the presence of pyruvate, α-alanine, and L-glutamine could consistently be substituted with D-glucose without detrimental impacts to cell growth ^44^. This finding was later corroborated by Eagle *et al*., who noted that, in contrast to D-glucose-based formulations, only a small fraction of the D-galactose metabolised by cells was converted to lactic acid, thereby preventing acidification of the medium ^45^.

This consideration is particularly critical, as human muscle cells exhibit a propensity for heightened glycolytic activity when cultured in supraphysiological glucose concentrations, compared to muscle tissue *in vivo* ^46^, a phenomenon known as the Crabtree effect ^47^. The use of D-galactose and L-glutamine as primary carbon sources forces cells to have an increased reliance on oxidative phosphorylation for ATP production due to negligible net ATP gain from the metabolism of D-galactose ^48,49^. The resultant metabolic shift serves additional purposes. Firstly, it renders the cells more susceptible to mitochondrial insult by attenuating compensatory glycolysis ^50^. Secondly, it enhances the comparability of the constructs to the metabolic profile of native type I (slow oxidative) muscle fibres.

In scenarios where specific medium formulations are required to sustain cell proliferation or induce differentiation, and a CO₂-free system is not viable, flight hardware can be ‘flooded’ with a defined gas mixture tailored to the needs of the cell type. However, this approach was unsuitable for the MicroAge experiment, as the flight hardware was vented to facilitate gas exchange with the external environment. Alternatively, incubation facilities that have an in-built gas supply are also available ^51^.

### Selection of an Upload Temperature Range for the Muscle Constructs

During the mission campaign, the launch and upload phases were limited by the absence of active thermal regulation and power availability in the launch vehicle, thereby preventing autonomous medium exchanges in the EUs. The lack of medium refreshes was anticipated to result in nutrient depletion and the accumulation of metabolic waste products, potentially compromising the viability of the muscle constructs inside the limited volume of the CC.

The basal metabolic rate of mammalian cells is regulated by a complex interplay of intrinsic and extrinsic factors, with temperature serving as a critical determinant ^52^. The potential to reduce the metabolic rate of AB1167 myotubes, by lowering the ambient upload temperature was investigated, and this approach aimed to reduce nutrient consumption and preserve cell viability during an extended stowage period (**Fig. 10**) ^53^.

The findings illustrated in **Fig. 10** demonstrate that AB1167 myotubes are capable of withstanding sub-physiological culture temperatures (ranging from 22-30 °C) in Leibovitz L15 medium for up to 120 hours without the necessity for medium exchanges. Further, the viability results were comparable with those observed with the DMEM-based control. Consequently, thermal bricks preconditioned to +30 °C were selected for upload, as they were reported by NASA to maintain Double Cold Bag (DCB) temperatures between +30 °C and +25 °C for up to 200 hours.

While cellular responses to sub-physiological culture temperatures remain incompletely understood, it is generally acknowledged that such conditions induce cell-cycle arrest and diminished proliferation (where applicable), alterations in mRNA transcription and translation, a reduction in metabolic activity, decreased accumulation of toxic by-products (e.g., lactate), and an overall downregulation of cellular processes ^54^. However, the specific temperature range investigated in this study may not be universally suitable for all mammalian cells and experimental parameters should be defined for specific cell types ^55^.

### Evaluation of the Impedance Sensing System to Monitor Muscle Contractions

Investigating the effect of electrical stimulation on muscle construct responses was a key objective of the project. However, the hardware’s spatial limitations prevented the inclusion of force transducers or an optical system to verify construct contractions after electrical stimulation. Attenuated responses to contractile activity have been suggested as a potential mechanism for muscle atrophy in microgravity, making it essential to confirm that contractions have occurred ^56^. It was proposed that geometric alterations of the myotubes within the muscle constructs, both during and following contraction, would result in a detectable change in the impedance of the complex biological material ^57–59^.

The system used platinum electrodes to introduce a low intensity ∼200 mV AC signal between stimulations, enabling impedance measurement. The data collected (**Fig. 11**) demonstrated that the impedance spectrum could effectively differentiate between empty scaffolds, non-viable constructs, and actively contracting constructs. As a result, this method was adopted to confirm that contractile activity occurred in the electrically stimulated muscle constructs during the mission.

However, a drawback of the system was that the impedance data represented an average across the entire scaffold, encompassing all three muscle constructs, rather than providing measurements at the level of the individual construct. As a result, it was not feasible to assess the change in impedance of individual muscle constructs within each separate (CC), and therefore, it was not possible to empirically confirm that each muscle construct contracted equivalently.

Lastly, the power limitations of the Kubik incubator rendered the concurrent execution of the electrical stimulation protocol across all relevant units unfeasible (**Table 2**). As such, the protocol execution was conducted in a staggered manner.

### Experimental Design Limitations

The mission was designed to maximise the utilisation of the Kubik incubator, enabling the accommodation of 72 skeletal muscle constructs on board the ISS. The muscle constructs were arranged three per scaffold across 24 experiment units, providing a minimum of six constructs per experimental group (**Table 2**). This design strategy was implemented to address a common limitation in microgravity research; the typically low number of biological replicates feasible during orbital missions and/or limited flight opportunities for repeated experimentation ^60^. This point also applies to studies of human muscle from astronauts in microgravity, due to the limited number of crew members available on the ISS from which samples can be obtained, in combination with limited public data availability ^61^.

While efforts were made to increase replicate numbers, it should be noted that each experimental group (6 constructs) was distributed across only two EUs, with constructs within an EU sharing a common scaffold, culture medium source, and electrical stimulus where applicable. As such, these constitute pseudo-replicates, rather than true independent biological replicates, and this limitation must be considered when interpreting the mission data.

A further limitation relates to the AB1167 cell line, which was originally derived from a 20-year-old male donor ^19,20,35^. Consequently, the study utilises cells from a single source, restricting the ability to capture inter-individual variation in responses to microgravity, HSP10 overexpression, or electrical stimulation

Lastly, the ground reference experiment (GRE) was conducted post-flight, following the return of samples from microgravity. This approach was necessary due to the absence of real-time environmental data downlink (e.g. temperature). Environmental data first had to be retrieved from onboard data loggers (iButtons) before the mission could be accurately simulated on the ground. Implementing real-time data downlink would enable parallel execution of the GRE and spaceflight experiment.

### Conclusions

These preparative studies were completed, and the MicroAge experiment was subsequently launched to the ISS on board the SpaceX 24 cargo resupply mission (CRS) on 21 December 2021 where it was installed into the Kubik incubator onboard the Columbus module. The fixed and frozen samples were returned to earth ∼1 month later.

All data suggested that the hardware functioned nominally, with novel impedance measurements and recovered data, along with biological samples, confirming the model’s suitability for studying the biological effects of microgravity on human skeletal muscle constructs. These findings are currently being prepared for further publication.

## 6 Author Contributions

Author SWJ prepared the article. Authors SWJ, SS, BT, DT and KH contributed to the generation and analysis of data and figure preparation. Authors GN, WB, GO, CM, AJ and KH were responsible for hardware development activities. Authors KH, FM, AJ, CM and JH and contributed equally to their critical appraisal of the manuscript. Authors MJJ and AM conceptualised the experiment and edited the manuscript. All authors approved the final submitted article.

## 7 Acknowledgements

The human skeletal muscle cell line AB1167, derived from the *Fascia lata* of a 20-year-old male, was utilised in this study and supplied by the Myoline platform at the Institut de Myologie, Paris, France ^19^.

---

**Supplementary Figure 1** Video of a AB1167_CTL muscle constructs contracting during the electrical stimulation protocol on ground.

